# Analyzing Human Movements - Introducing A Framework To Extract And Evaluate Biomechanical Data

**DOI:** 10.1101/442087

**Authors:** Chris Richter, Enda King, Siobhan Strike, Andrew Franklyn-Miller

## Abstract

This study discusses possible sources of discrepancy between findings of previous human motion studies and presents a framework that seeks to address these issues. Motion analysis systems are widely employed to identify movement deficiencies - e.g. patterns that potentially increase the risk of injury or inhibit performance. However, findings across studies are often conflicting in respect to what a movement deficiency is or the magnitude of the relationship to a specific injury. To test the information content of movement data, a framework was build to differentiate between movements performed by a control (NORM) and abnormal (IMP-L and IMP-C) cohort using solely movement data. Movement data was recorded during jumping, hopping and change of direction exercises and was mathematically decomposed into subject scores. Subjects scores were then used to identify the most appropriate machine learning technique, which was subsequently utilized to create a prediction model that classified if a movement was performed by: IMP-L, IMP-C or NORM. The Monte Carlo method was used to obtain a measure of expected accuracy for each step within the analysis. Findings demonstrate that even the worst classification model outperformed the best guess observed and that not all members of the NORM group represent a NORM pattern as they were repeatedly classified as IMP-L or IMP-C. This highlights that some NORM limbs share movement characteristics with the abnormal group and consequently should not be considered when describing NORM.

## Introduction

Motion analysis systems are widely employed within both universities and clinical facilities,to explore the association between movement patterns and athletic performance and possible risk of injury, which is of great interest to coaches, physiotherapists and other medical professions. However, while there are many studies that have examined movements using kinematic and kinetic measurements [1,2] seeking to identify features, such as maximum knee flexion, that might be related to injury, there are no well established evidence-based guidelines that state what a movement deficiencies is, what normative ranges of variability are or what a “normal” movement looks like. Additionally, there is little agreement as to which, if any, movement tasks (exercises) can expose patterns that could lead to injury or what properties a exercise should hold (single or double leg, movement in one, two or three planes, ecological valid and so on). For example, both Hewett et al., [1] and Krosshaug et al., [2] examined features extracted from a double leg drop jump (DLDJ) to assess their ability to predict the risk of sustaining an anterior cruciate ligament (ACL) injury. Both studies performed a prospective analysis on a large population of female athletes participating in field sports. While Hewett et al., [1] concluded that features (describing knee motion and knee loading) were very sensitive in predicting ACL injury, Krosshaug et al., [2] was not able to predict ACL injury using features extracted from the DLDJ. With such conflicting results between studies it is no surprise that, in a recent publication, Bahr [3] came to the following conclusion: “To date, there is no screening test available to predict sports injuries with adequate test properties and no intervention study providing evidence in support for screening for injury risk.” However, while predicting acute injuries from movement features is challenging (as there may be multiple contributing factors to injury risk - e.g. movement deficiencies, fatigue, contact with other players, genetics, environmental factors, changes in acute/chronic training load, lost focus, frequency of movement assessments), studies examined within Bahr’s [3] review utilized features independently (e.g. peak knee valgus moment, maximum knee flexion on their own), rather than the interaction of features. Further, to date, there has been little progress in the way biomechanical features are extracted and analyzed and this could be a reason for the disparity between studies. Traditional analyses often discard a large portion of the captured data [4–7] and do not consider the interrelationships within such a complex and multivariate system as the human body.

When describing a movement, studies often extract features based on prior knowledge (previous research and / or personal and clinical experience) or post hoc analysis, performing a comparison of magnitude or timing [8] assuming that these features capture the underlying function of a signal. While discrete points can be helpful in understanding movements, the selection of discrete points has the potential to discard important information [4,5], to compare features that present unrelated neuromuscular capacities [6] and to encourage fishing for significance - e.g. non-trivially biased non-directed hypothesis testing [7]. Due to the apparent limitations in discrete point selection, other analyses have been introduced in recent years - statistical parametric mapping [7], (functional) principal component analysis [9–11], analysis of characterizing phases [6], point by point manner testing[12] and other techniques [8,13,14] to improve the analysis of movement patterns.

Another possible source for conflicting conclusions is the way extracted features are compared across groups. Comparisons are often made using statistical significance, which bases conclusions by inferring properties about a population by testing hypotheses and deriving estimates by probability values (p-values). While, p values are useful they have been criticized in the past and present [15–19]. Criticism of p values include the possibility of committing type 1 and 2 errors, that any difference can be statistically significant - with large enough sample size [18] - that p values do not provide statistical precision and that conclusions do not account for subgroups within the data. The way an athlete moves could be influenced by anthropometric measures, sporting and injury history and there is growing evidence for differences within movement strategies across individuals [20–29]. As such, features that relate to the risk of injury could be masked during an analysis because of different movement strategies.

Finally, most movement analysis studies involve the recording of multiple trials but commonly examine the average of all or the maximum trial. When averaging multiple trials, an ?artificial? movement is created and examined. Local peaks will be altered in magnitude and temporal appearance [30] and the intersegmental link (coordination) between joints might be lost. As such, examining the best trial seems more valid. However, such an approach selects a unique instance (based on jump height or performance time) and could bias an analysis towards a non-realistic situation. No athlete will perform a task over and over with a maximal effort and consequently the sub-maximal efforts should not be discarded. An alternative approach is to utilize repeated random sampling (Monte Carlo simulation), where the captured trials are selected at random and the analysis is run multiple times [31,32]. This can overcome the “maximal effort bias” and can also provide a measure of expected differences or accuracy within future studies. Further, such an approach can also overcome discrepancies between findings that are caused by the selection of a reference limb when comparing a abnormal (injured) to a normal or uninjured group.

When examining a human movement, the method of data analysis chosen needs to contend with a complex and multivariate system and any analysis using an inference test may not help to progress the understanding of movement further as it does not account for differences in movement strategy or the interrelationship of segments. More suitable methods for movement analysis might be machine learning techniques, which have gained popularity in other fields and have demonstrated an enhanced ability to understand complex and multivariate system - see Rajpurkar et al., [33] or Barton et al., [34]. Rajpurkar et al., [33] demonstrated the use of such an approach in cardiovascular medicine, by developing an algorithm that outperformed board certified cardiologists when detecting a range of heart arrhythmias from electrocardiograms. When applying machine learning techniques to motion analysis, the computer is trained to learn the underlying connection between a set of predictor features (e.g. peak knee angle and moment) and a class (abnormal or normal). If the set of predictor features hold sufficient information, the algorithm will be able to predict the class of previously “unseen” observation correctly. If the set of predictor features does not hold sufficient information, the algorithm will fail to predict the correct class of a previously “unseen” observation. To build a model that can objectively judge a movement, Richter et al., [29] proposed identifying individuals that present the “true” group pattern - e.g. are continuously classified into the correct class. For example, if the goal is to describe a normal movement pattern, an algorithm could be trained to differentiate between a normal and abnormal population. Samples, which cannot be correctly classified by the classification technique, might not be considered when describing the normal behavior. Consequently, the probability of belonging to a class (abnormal or normal) generated during a classification could be used when judging a movement [29]. A good example of an abnormal cohort are athletes recovering from ACL reconstruction as the ACL reconstruction will have altered the neuromuscular properties of the athlete, influencing the way they move.

The aim of this study was to examine if biomechanical data can distinguish between a normal and abnormal movement pattern and to present a framework that combines an automatic feature extraction with a machine learning approach.

## Materials and methods

### Subjects

The data set used in this study holds a cohort of athletes recovering from an ACL reconstruction and an uninjured control group. The ACL group (n = 156) was recruited from the caseload of two orthopedic surgeons who specialize in knee surgery between January 2014 and December 2016. Inclusion criteria were: biomechanical assessment approximately 9 months after ACL reconstruction, intention of returning to full participation in multi-directional sport after surgery, bone patellar tendon bone graft or hamstring graft from the ipsi-lateral side, gender (male) and an age between 18 and 35. Exclusion criteria were: multiple or previous ACL reconstructions and any meniscal repairs. The control group (n = 62) contained males that participated in multi-directional sport (i.e. Gaelic Football, Soccer, Hurling, Rugby Union) that were free of injury in the 3 months prior to testing, had no previous knee surgery and were between 18 and 35 years of age. The study received ethical approval from and was registered on clinicaltrials.gov.

The ACL group had an average age of 24.8 ± 4.8 years was 180 ± 8 cm tall and had a body mass of 84 ± 15.2 kg. The control group had an average age of 24.8 ± 4.2 years was 183 ± 6 cm tall and had a body mass of 82 ± 8.9 kg.

### Data Capture and Preprocessing

The testing took place in the motion analysis laboratory using an eight-camera motion analysis system (200Hz; Bonita-B10, Vicon, UK), synchronized with two force platforms (1000Hz BP400600, AMTI, USA). Before data collection, all subjects undertook a standardized warm-up and wore their own athletic footwear with 24 reflective markers secured to the shoe or to the skin using tape, at bony landmarks according to the Plug-in-Gait marker set. Three trials of each limb for the following exercises were captured: DLCMJ [35], SLCMJ [35], DLDJ [35], SLDJ [35], HoHo [35], SLHop [35], CoDP [36] and CoDU [36]. Marker and force data were low-pass filtered using a fourth-order Butterworth filter [37] - before computing kinematic and kinetic measures using Nexus (1.8.5; Vicon, UK). All kinetic variables were normalized to body mass. Data pre-processing (gap filling and waveform screening) was performed using a custom developed MATLAB program (R2015a, MathWorks Inc., USA) that also computed the additional kinematic measures [35]. The start and end of an exercise was defined using the force trace or a combination of center of mass (CoM) power and force trace. For the CMJ the start was defined as the first time the ground reaction force (GRF) was less than BW-25N, while the end was defined as toe off (force less than 25). For the DJs, HH and CoDP the start was defined as the first instance the vGRF is above 25N and the end when the GRF is below 25N. For the SLHop the start was defined, as the first instance were GRF was above 25N and the end when the center of mass (CoM) power first became positive. All measures were landmark registered using a dynamic time warping process [38] to align the end of the eccentric phase across the all curves^1^.

### The Framework

The steps taken during data analysis can be described as follows: feature generation, selection of a supervised learning technique, and generation of a classification model and testing of the generated classification model. During the analysis, each limb was treated as a separate entry to overcome the question of which limb to choose within the control group. Consequently, the data that was used during the following steps contained 156 operated limbs (class labeled as IMP-L), 156 non-operated limbs (class labeled as IMP-C) and 124 control limbs (class labeled as NORM). The description of the generation of the framework is based on a single exercise and was done for every exercise separately. Every process described was performed 100 times using different randomly chosen trials to obtain a robust measure of findings. A flowchart of the process and aim of each step is illustrated in fig 1. Within this study, only the maximum trial of the 3 captured (based in jump height, contact time) was chosen for data analysis.

**Fig 1.**
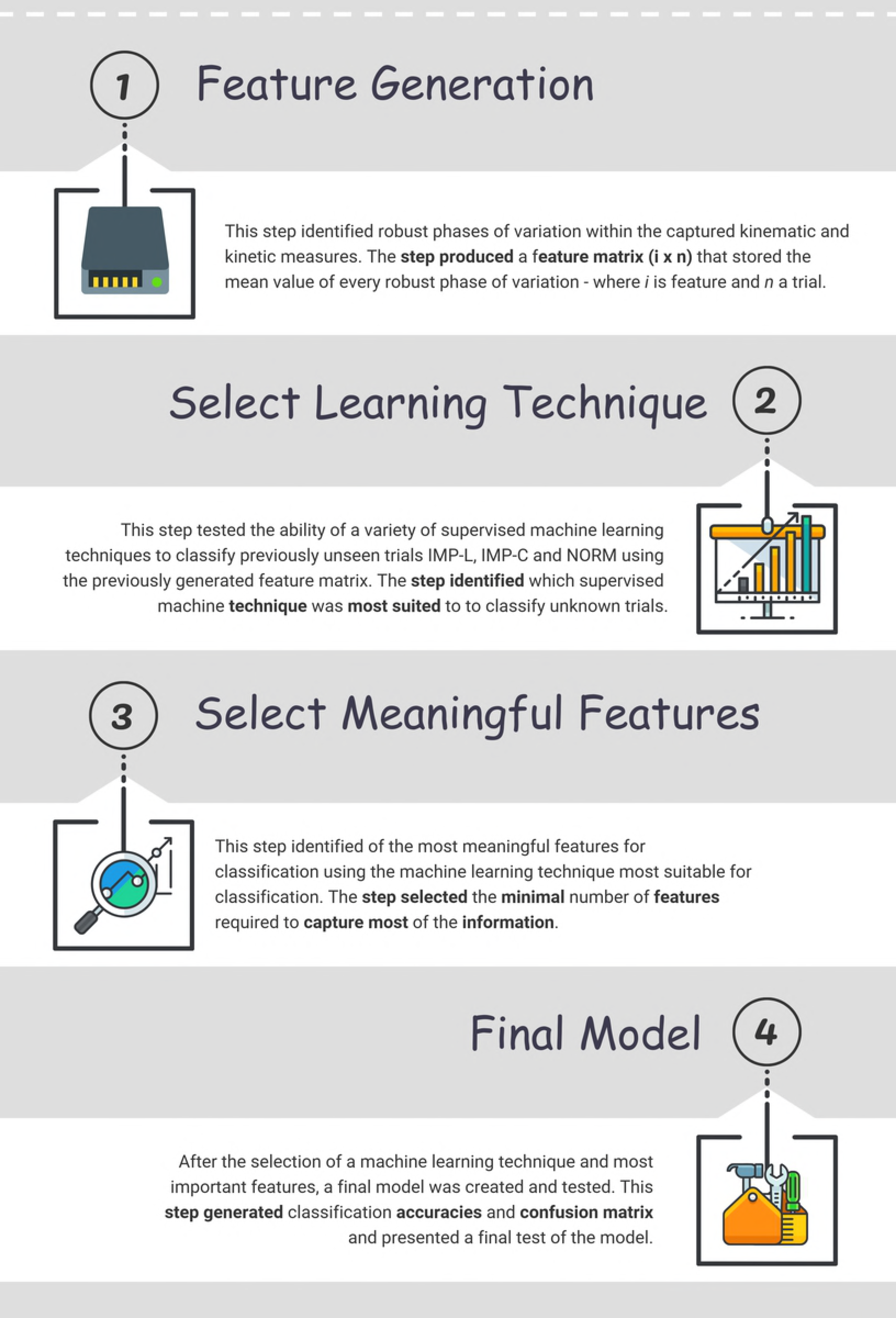
Illustration of the four steps performed during the generation of the classification model.

### Feature Generation

The first step was the identification of phases of variation, which were used to calculate features that describe the behavior of a trial - similar to Richter et al., [39]. The process is illustrated in detail in fig 2.

**Fig 2.**
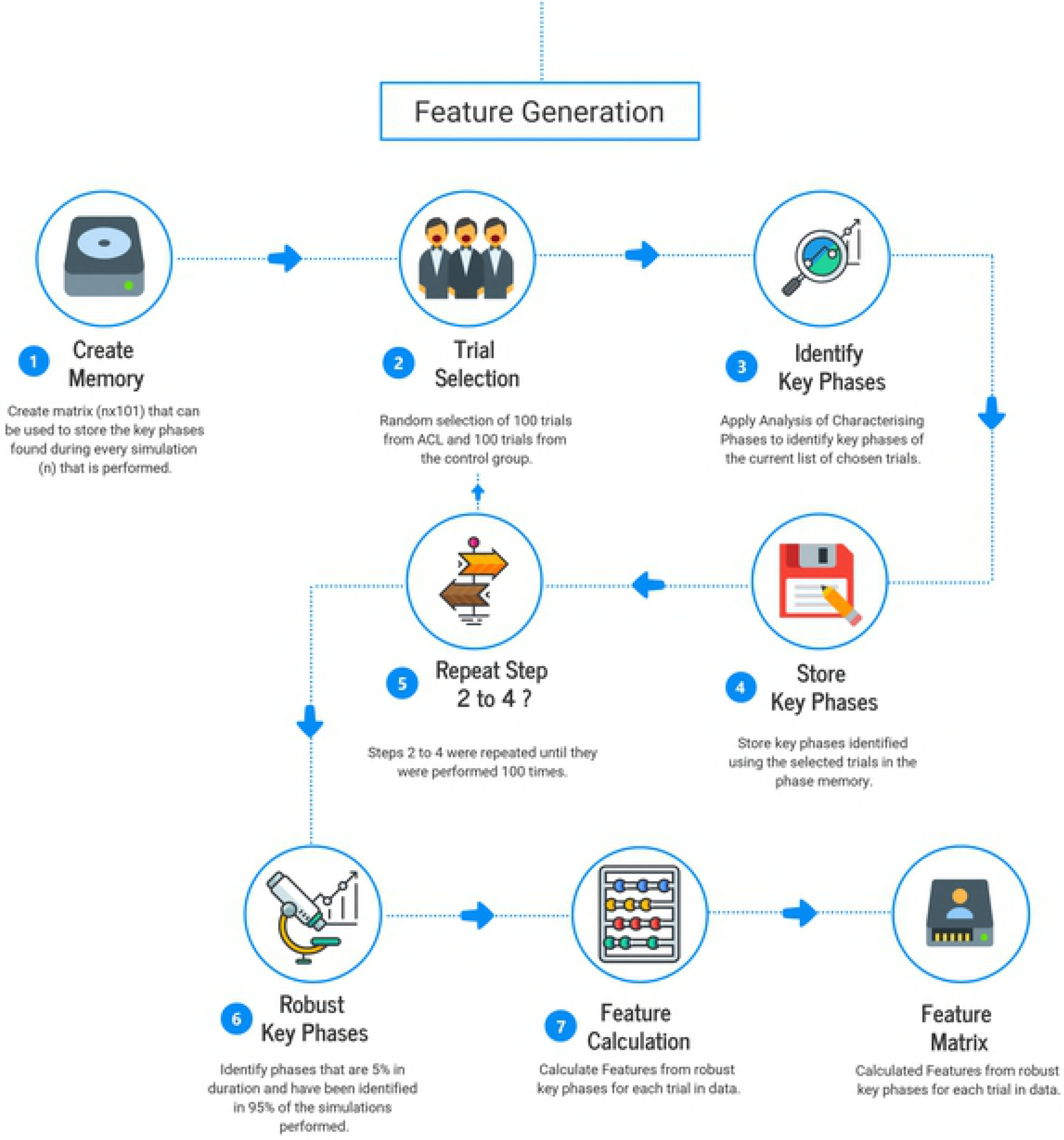
Illustration of the steps performed to generate a features matrix that contains the calculated subject scores.

To identify phase of variation, 100 trials were chosen randomly from the IMP-L and IMP-C class as well as 100 trials of the NORM class, forming a dataset of 200 trials. This dataset included a variety of measures^2^ and phases of variation where identified using the idea of analysis of characterizing phases [6]. The information obtained (measure [e.g. joint angle], start and end) of each phase of variation found was recorded. This process was repeated 100 times, omitting the not selected trials (n = 236) during each simulation to increase the generalizability of findings. All phases that occurred at least 95 times, during the 100 iterations, and spanned over at least 5 % of the measure were considered “robust” and used to generate a feature matrix (i features x 436 trials) to describe the movement pattern within a trial. A feature i was calculated as the average value over a robust phase of variation. For every double leg exercise, symmetry was also calculated as limb - contralateral limb - e.g. left - right and right - left.

### Selection of Supervised Learning Technique

The second step was the identification of the most appropriate machine learning technique, as every technique has different abilities to learn relationships between a classes (IMP-L, IMP-C and NORM) and features that are used to train the algorithm [40,41]. The process is illustrated in detail in fig 3 and was run twice: once using raw features and ones using normalized features (z-scores).

**Fig 3.**
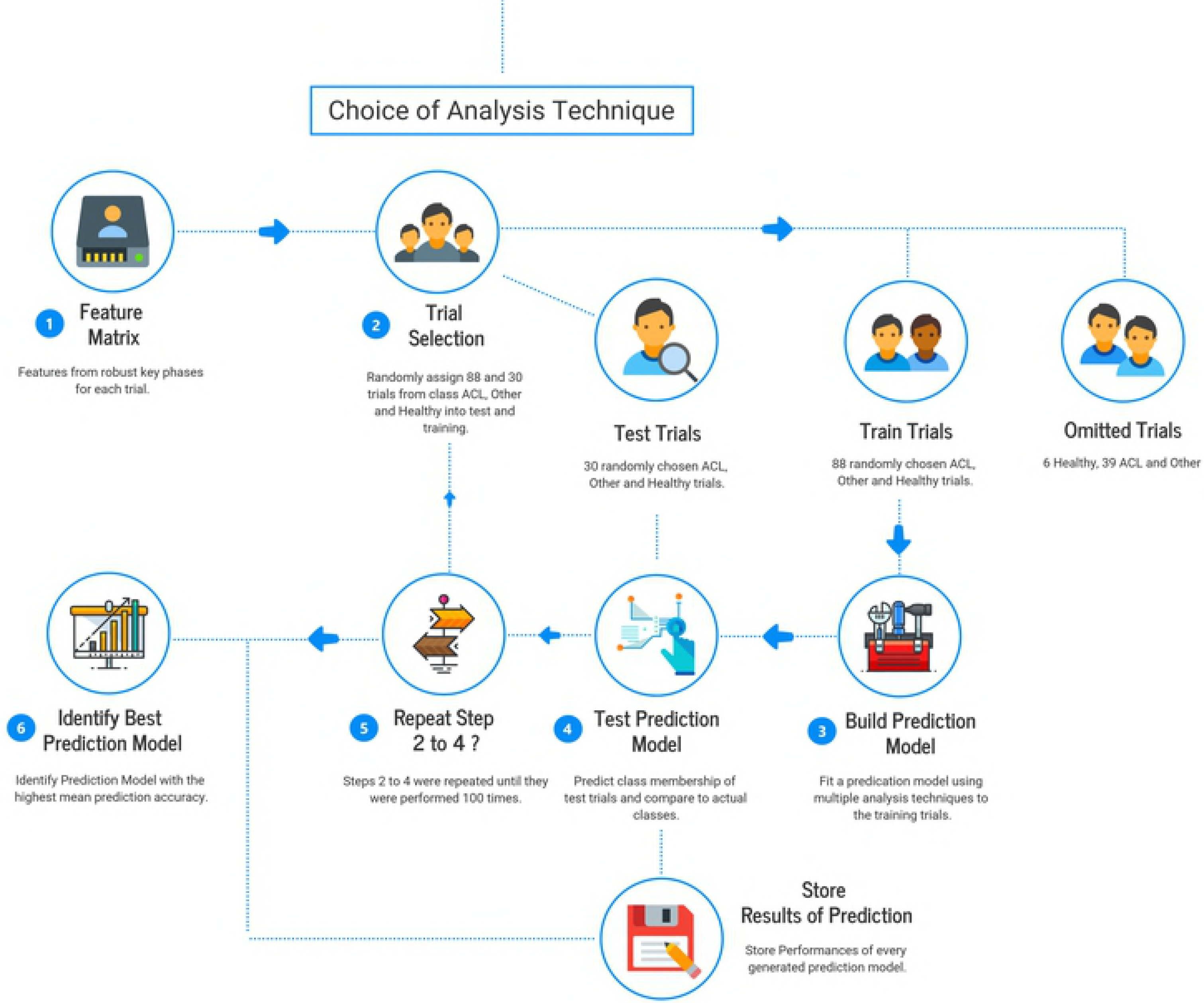
Illustration of the steps performed to identify the most suitable learning technique.

This study examined the ability of techniques included within the statistical toolbox of Matlab (R2015a, MathWorks Inc., USA): a decision tree (fitctree), an ensemble of decision trees (n trees = 50; TreeBagger), a discriminant analysis model (fitcdiscr), a naive Bayesian classifier (fitcnb), a k-nearest-neighbor model (fitcknn), a multi class model for support vector machines (fitcecoc), a regression model (mnrfit; in stepwise forward) and a neural network (patternnet). To identify the best learning technique for the classification of the classes (IMP-L, IMP-C and NORM), the data was split into training, test and leave out dataset. The sample size of each data set was chosen based on the NORM sample size (train = 70 %, test = 25 % and leave out = 5 %). Hence, the train dataset contained 88 trials from each class (about 56 % of IMP-L and IMP-C) resulting in n = 264. The test dataset contained 30 trials from each class (about 20 % of IMP-L and IMP-C) resulting in n = 90, while the leave out dataset contained the remaining trials - 6 NORM, 38 IMP-L and 38 IMP-C (about 24 %). The train set was used to teach the examined machine learning techniques to forecast the three classes based on the extracted features. The purpose of the leave out data set was the “interchange” of trials within simulations to increase variation across data sets. After the training had been completed, comparing the predicted class to the actual class of the test data assessed the performance of each learning technique. The accuracy of each model was recorded and the process was repeated 100 times using different randomly selected training and testing samples. The learning techniques with the highest mean accuracy over the 100 repetitions was judged most appropriate.

### Selection of Features

The third step was the identification of the most meaningful features for classification. This process used only the best performing machine-learning technique and sought to reduce the effect of over-fitted models by identifying the minimal number of features required to capture most of the information within an exercise. The process is illustrated in detail in fig 4.

**Fig 4.**
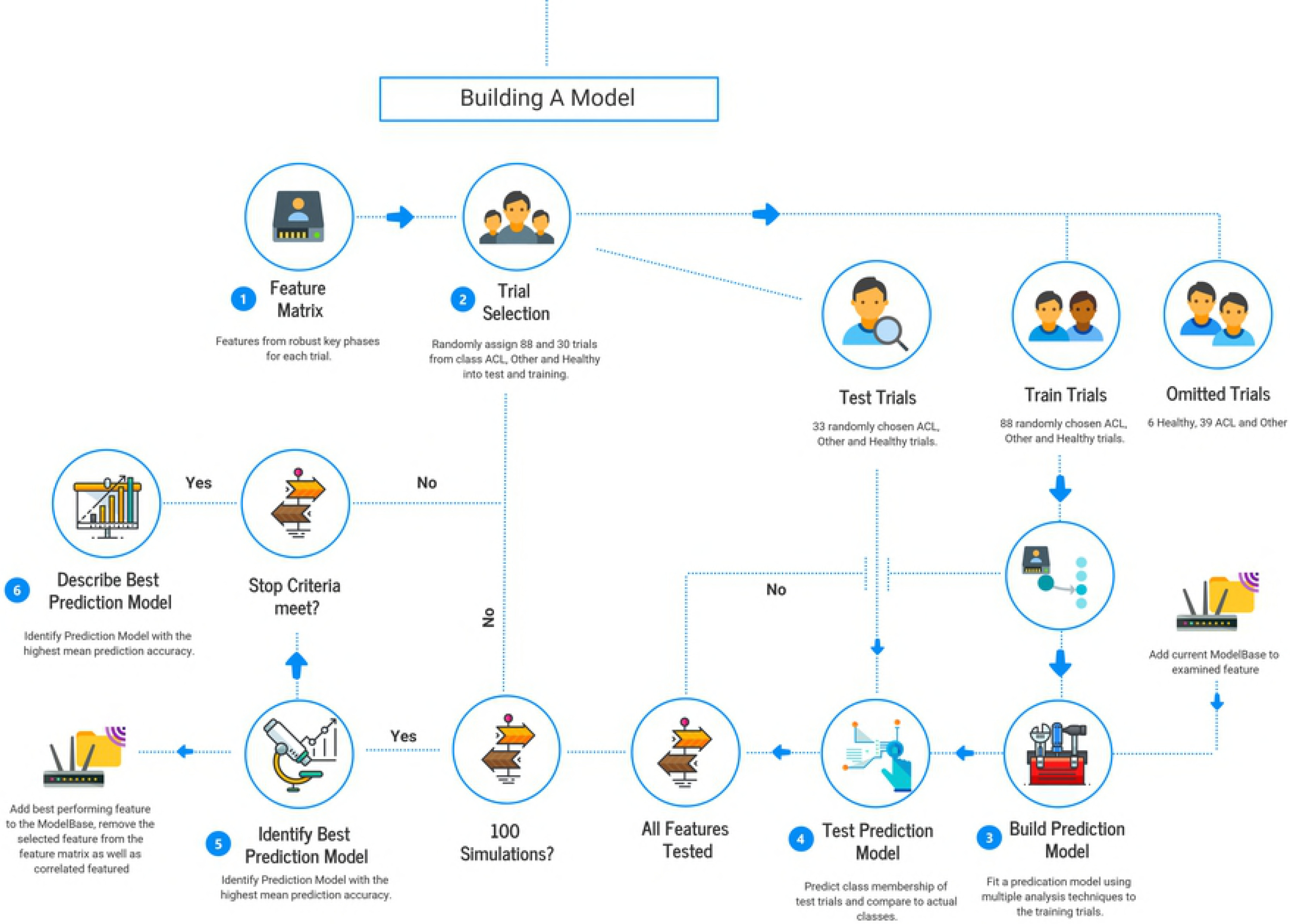
Illustration of the steps performed to identify key features toward classification and to generate the to be tested classification model.

To identify features with high importance towards classification, the data was, as before, split into train, test and leave out data sets (see selection of supervised learning technique). The train set was used to teach the best performing machine-learning technique to forecast the three classes (IMP-L, IMP-C and NORM). After the training phase had been completed, the performance of the learning technique was compared to the predicted class of the test data to the actual class. This process was done for each extracted feature on its own. The accuracy of each “feature model” was recorded and the process was repeated 100 times using different randomly selected train, test and leave out samples to obtain a repeatable measure of the expected accuracy.

Subsequently, the feature with the highest mean accuracy was identified, removed from the feature matrix and used to build the “model base”. All features that correlated with the identified feature (greater than 0.7) were removed to increase interpretability based on a correlation utilizing the whole feature matrix. After the model base was built, the process was repeated while pairing every feature remaining in the feature matrix with the model base. The feature performing best in combination with the model base was added to the model base, multi-collinear features were removed. This process was repeated until 20 features were added to the model base (this number as chosen to reduce computing time). To find the “optimal” prediction model, the model with the smallest number of features that accounts for a large part of the maximal observed accuracy, the Elbow method was used - described in Hastie and Tibshirani [40] or Vapnik and Vapnik [41]. The “elbow” was defined as the point n where the differentiation of accuracy f improved less than 10 % of its range (Eq 1).

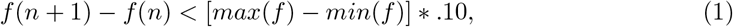

### Generation and Test of the Final Classification Model

After the selection of a machine learning technique and most important features, a final model was created and tested. As before, the data was split into train, test and leave out data sets. The best performing machine-learning technique was trained to predict classes? using the optimal number of features and comparing predicted and actual class of the test data assessed prediction performance. During each simulation, false and correct classifications of each trial were recorded to allow the generation of a confusion matrix and to examine the consistency of true classifications for each trial. This allowed the examination if a generated model had difficulties to differentiate classes, e.g. NORM can be separated from IMP but IMP-L and IMP-C cannot be separated from each other (demonstrate the same pattern), and helps to understand and interpret the findings. During the simulations, class allocation by guessing were also performed for every trial within the test data, with each trial having a 1 in 3 trial chance of belonging to the class: IMP-L, IMP-C or NORM. A classification model was defined meaningful if the lowest observed accuracy during the iteration exceeded the best-observed accuracy during guessing. Matlab (R2015a, MathWorks Inc., USA) was used for data processing and analysis.

## Results

### Feature matrix and sample selection

The number of identified phases of variation differed between the examined exercises (78 to 232). During the key phase selection, trials from the IMP-L and IMP-C class were selected on average 35 times (min 21 - max 50), while trials from the NORM class were selected 85 times (min 76 - max 92). During the model generation, IMP-L trials were selected on average 48 times (46 to 49) for the training process and 19 times (11 to 34) for the testing process. Each IMP-C trial was selected on average 47 times (46 to 50) to train and 19 times (11 to 34) to test a classification model, while each NORM trial was selected on average 75 times (63 to 85) to train and 25 times (15 to 37) test a classification model.

Reported findings are based on normalized scores (z-scores), as the difference between classifications accuracies from raw and z-scores were at a maximum of 1 % and because z-scores remove any possible magnitude effect during the selection of features for the final model (table 1).

**Table 1.** Description of performances, differences between conditions observed during each step of the data analysis as well as performances for a 2 condition classifier (IMP and NORM) and percentage to NORM with “true” group membership.

| Exercise | Accuracy best classifier (all features) | Difference z-scores vs. raw | Accuracy best classifier (20 features) | No. optimal features | Accuracy best classifier (optimal features) | % with only IMP & Norm class | % Norm presenting true pattern |
| --- | --- | --- | --- | --- | --- | --- | --- |
| DLCMJ | 73% | 0 | 80% | 4 | 74% | 78% | 60% |
| SLCMJ | 78% | 0 | 80% | 2 | 78% | 99% | 99% |
| DLDJ | 82% | 0 | 88% | 4 | 82% | 83% | 50% |
| SLDJ | 66% | 0 | 66% | 5 | 64% | 73% | 42% |
| SLHop | 75% | 1 | 79% | 4 | 72% | 87% | 75% |
| HuHo | 67% | 1 | 71% | 3 *15 | 60% *70% | 74% *85% | 50% |
| CoDP | 66% | 1 | 72% | 5 *18 | 58% *72% | 72% *86% | 58% |
| CoDU | 66% | 0 | 66% | 2 *18 | 51% *67% | 67% *78% | 35% |

\* findings of manual selection of the number of optimal features

### Best performing machine-learning techniques

The ability to classify an unseen trial into IMP-L, IMP-C and NORM using every extracted feature, differed between the machine learning techniques within and across exercises. Their highest accuracy for each exercise, in decreasing order, was as follows (machine learning technique; mean; range; 95 % confidence interval): double leg (DJ) drop jump (DJ; RandomForest; 82 %; 71 to 91 %; 81 to 83 %), single leg (SL) countermovement jump (CMJ; Regression; 78 %; 68 to 87 %; 77 to 79 %), SLHop (Regression; 75 %; 64 to 86 %; 74 to 76 %), DLCMJ (Regression; 73 %; 62 to 85 %; 72 to 74 %), Hurdle Hop (HuHo; Discriminant; 67 %; 56 to 77 %; 66 to 68 %), planned change of direction (CoDP; Discriminant; 67 %; 54 to 77 %; 66 to 68 %), SLDJ (Discriminant; 66 %; 54 to 78 %; 65 to 67 %), unplanned change of direction (CoDU; Discriminant; 66 %; 48 to 79 %; 65 to 67 %). A detailed illustration is given in table 1. For some exercises (DLDJ, SLCMJ, HuHo and CoDP), the performances of many machine-learning techniques were very close to the best technique (fig 5).

**Fig 5.**
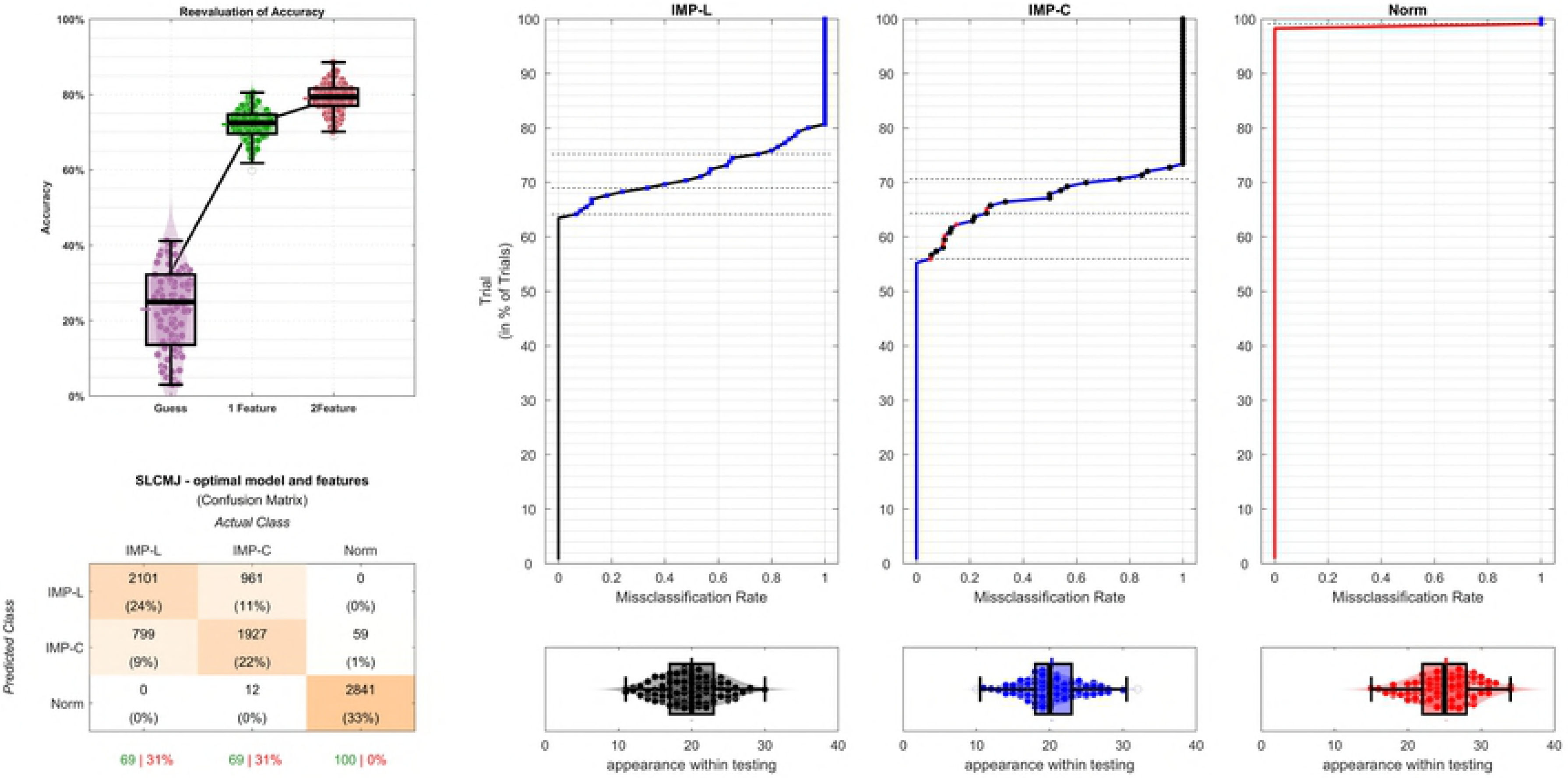
Performance of every learning technique within an exercise, using a z scores, illustrated by a violin plot overlaid by a box-plot. The shaded area represents the best performing learning technique.

### Performance classification model

Most models improved performance when using only 20 features, while using the “optimal” number of features generated classification accuracies slightly lower or equal (number of features; accuracy; difference to model using all features): DLDJ (4; 82; 0 %), SLCMJ (2; 78; 0 %), DLCMJ (4; 74; +1 %), SLHop (4; 72; -3 %), SLDJ (5; 64; -2 %), HuHo (3; 60; -7 %), CoDP (5; 58; -8 %) and CoDU (2; 51; -3 %). A detailed description is given in table 1. The best performing guess observed had an accuracy of 43 %.

## Discussion

### Model performances

This study introduced and examined the classification accuracy of a framework that combines an automatic feature extraction and a supervised machine learning technique, to differentiate movements performed by limbs with reconstructed ACL (IMP-L), limbs contralateral to an ACL limb (IMP-C) and limbs of a control group (NORM). Findings demonstrated that features extracted from biomechanical data, regardless of the examined exercise, contained enough information to outperform guessing by up to 40 % (based on mean performance and best observed guess within 100 simulations; 43 %). The best performing or most informative exercise was the DLDJ (82 %), which was followed by the SLCMJ (78 %), DLCMJ (74 %), SLHop (72 %), SLDJ (64 %), HuHo (60 %), CoDP (58 %) and CoDU (51 %). An effect of over-fitting to the training data was present in the models that used all extracted features, as modeling using a reduced number of features generally performed better at predicting the test data classes than models using every extracted feature. This highlights the needed for a cross-validation when applying a machine learning technique without using simulations. Within the optimal models (reduced number of features that captured most information) the number of features was reduced by 94 to 98 % but performances were similar to models using all features (approx. +2 %), except for models describing more complex movements (HuHo, CoDP and CoDU; approx. -10 %). This highlights that a small number of features can generally capture a large portion of relevant discriminating information. While the best performing learning technique was affected little by using raw features or their z-scores, most other tested techniques responded positively tonormalizing scores (increasing performances close to best performing technique).

The examined exercises can be split in three groups based on their patterns within findings. The exercises DLDJ, SLCMJ, DLCMJ and SLHop represent non-complex movement. The HuHo, CoDP and CoDU represent a complex movement, while the SLDJ represents a non-complex movement with within “group-limb” confusions.

The DLDJ, SLCMJ, DLCMJ and SLHop are exercises that enabled the most accurate classification performances (72 and 82 %) using up to 4 features. When examining the confusion matrix of the SLCMJ and SLHop it was noted that a large portion of the prediction error (approx. 18 %) was caused by confusions within the IMP group (IMP-L and IMP-C), while there were only very few confusions of IMP-L or IMP-C with NORM (< 5 %; see fig 6).

**Fig 6.**
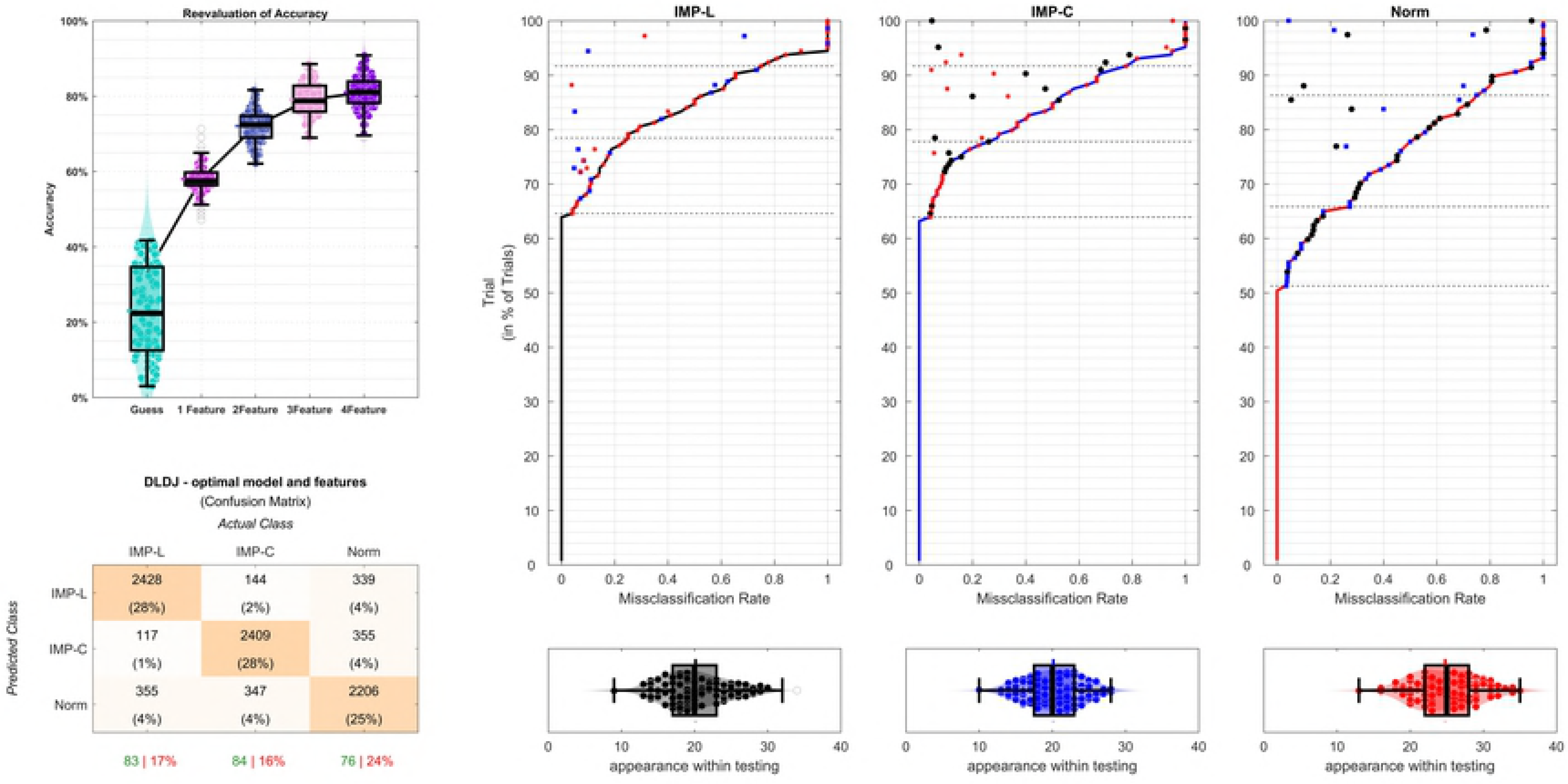
Illustration of the findings within the tested SLCMJ model. The model accuracy is displayed on the top right, while the confusion matrix of a 2-feature model is displayed below (right button). The three graphs on the left display the confusion pattern (button) and selection frequency of each trail within the IMP-L, IMP-C and NORM (top).

**Fig 7.**
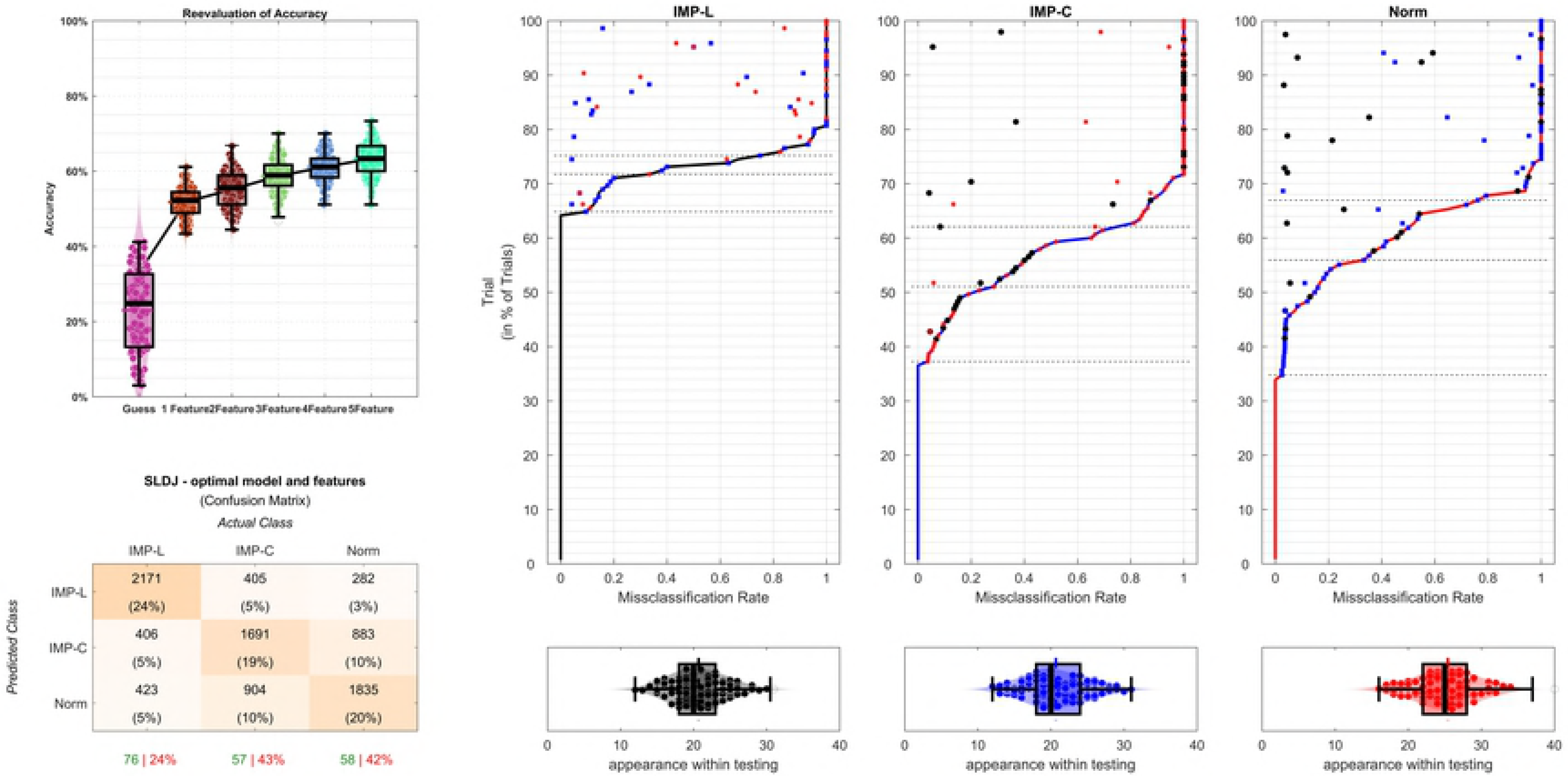
Illustration of the findings within the tested DLDJ model. The model accuracy is displayed on the top right, while the confusion matrix of a 4-feature model is displayed below (right button). The three graphs on the left display the confusion pattern (button) and selection frequency of each trail within the IMP-L, IMP-C and NORM (top)

If the classification accuracy had been assessed using only two classes (IMP vs. NORM), the classification accuracy of both exercises would have improved greatly - 99 % for the SLCMJ and 87 % for the SLHop. This might suggests that both limbs (IMP-L and IMP-C) are affected by ACL injury or in other words that both limbs deviate from the NORM pattern but not from each other. As such, the IMP-C may not be ideal to be used as reference when judging risk of injury. Another reason could be that the differences between IMP-L and IMP-C might be too small to be detected using the examined features and sample size. In contrast to the SLCMJ and SLDJ, the DLDJ and DLCMJ did not demonstrate an increased confusion pattern toward a specific confusion (e.g. IMP-C with IMP-L or IMP-C with NORM; see fig ??), which might be explained by the additional information: symmetry within the DLCMJ and DLDJ.

Symmetry features were selected first, followed by a performance approximate and another symmetry feature. The magnitude of symmetry or asymmetry seems to be a useful feature within these exercises and would support findings of Myer et al., [42]. While symmetry features could have been included within the single leg models, it would require the interventions of the investigators - e.g. symmetry calculation as mean symmetry, symmetry between trial x and trial y and so on [43]. As every exercise execution presents different external and internal conditions, symmetry cannot be calculated without setting subjective rules and was hence not included here. However, this also suggests that a testing battery should contain both single and double leg exercises.

The HuHo, CoDP and CoDU demonstrated a performance about 10 % less than DLDJ, SLCMJ, DLCMJ and SLHop within the optimal model. However, these exercises should not be discarded because the optimal model suffered from the detected number features - as they lost about 13 % of their prediction ability when the number of features was reduced from 20 to about 5. This suggests that these exercises are more complex and need more information to describe the underlying movement pattern. A reason could be that the initial condition within these exercises is less defined than in the DLDJ, SLCMJ, DLCMJ and SLHop. Jumping height, width and speed of the HuHo was not accounted for within the model nor had the model information (e.g. the feature jump height, width and so on) to adjust. Completion time and pre step movement pattern during the CoDP and CoDU was not accounted for within the model nor had the model information to adjust for these differences in execution. As such, there is a need for a larger number of features that is clearly demonstrated by an increased performance with the addition of features. Increasing the number of features to 15 in the HuHo resulted in a 10 % performance increase (70 %). Increasing the number of features to 18 in the CoDP resulted in a 14 % performance increase (72 %), while 18 features within the CoDU resulted in a 16 % performance boost (67 %). As with the SLCMJ and SLHop, a large portion of the error of the HuHo and CoDP originated from a “poor” ability to differentiate IMP-L and IMP-C. Assessing the classification accuracy for a two-class classification (IMP vs. NORM) resulted in a performance of 85 % for the HuHo (+ 15 %) and 86 % for the CoDP (+ 14 %). The prediction errors within the CoDU did not demonstrate an increased confusion pattern toward a specific confusion (e.g. IMP-C with IMP-L or IMP-C with NORM). A possible reason could be a combination of an increase in variability and physical abilities of subjects. The CoDU demonstrated the lowest percentage of always-correct classified NORM (only 35 % present the true pattern). These findings highlight that understanding a complex tasks requires many features in combination and that it becomes more challenging to extract a representative group movement with increasing complexity of a task.

The last remaining exercise was the SLDJ that demonstrated a rather “unusual” behavior. Its performance was comparable to the HuHo, CoDP and CoDU but its performance did not improve beyond the number of optimal features nor did it present the previously observed confusions pattern within the IMP group. However, a large portion of the prediction error (20 %) was caused by confusions between IMP-C and NORM, while there were fewer confusions of IMP-C or NORM with IMP-L (< 5 %, fig 8). If the classification accuracy had been reported for classifying only two classes ([IMP-C; NORM] and IMP-L) the performance of SLDJ would have improved to 83 %. As such the SLDJ seems to be able to extract movement patterns that describe if a limb had an ACL reconstruction or not.

**Fig 8.**
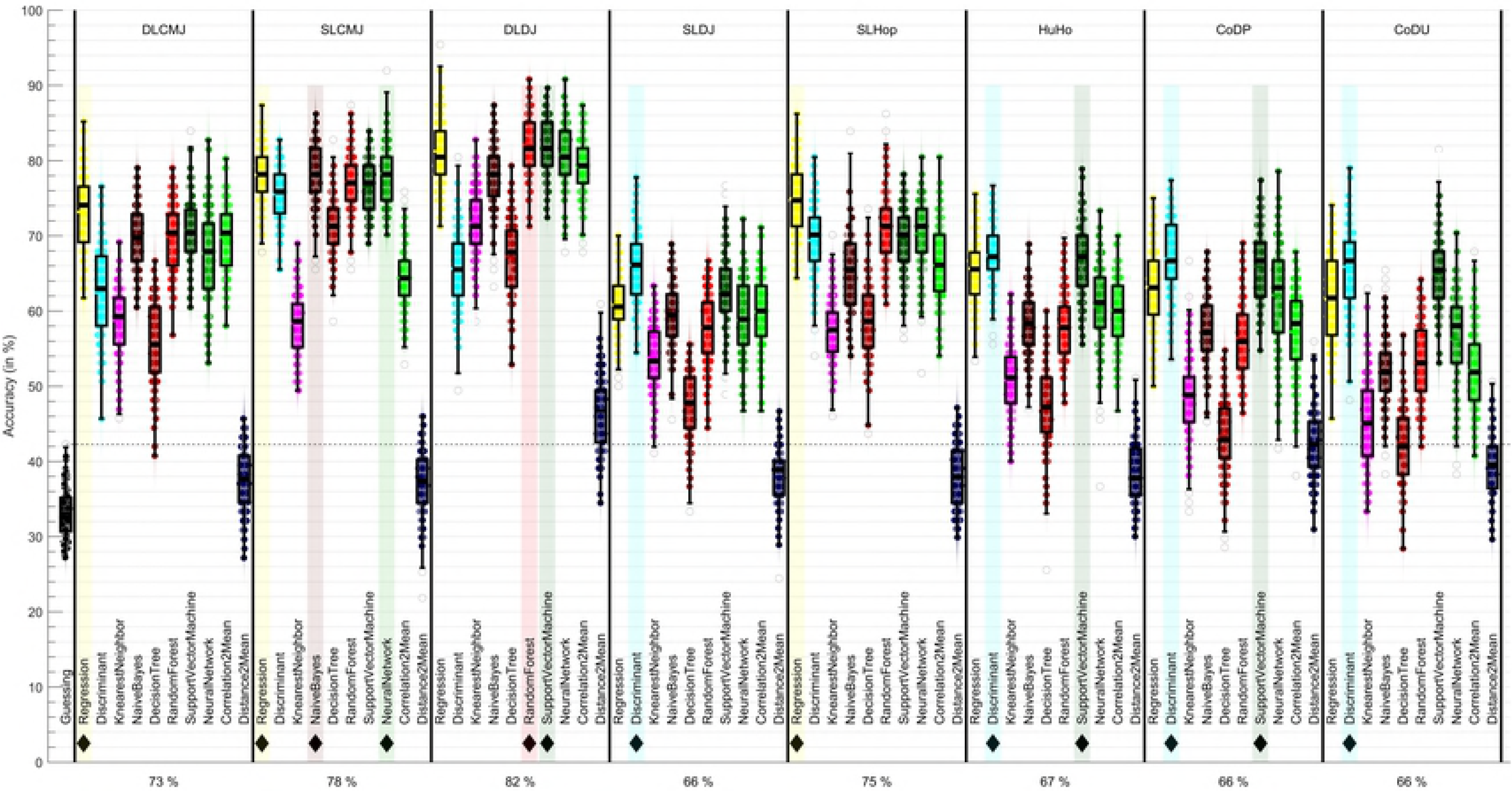
Illustration of the findings within the tested SLDJ model. The model accuracy is displayed on the top right, while the confusion matrix of a 4-feature model is displayed below (right button). The three graphs on the left display the confusion pattern (button) and selection frequency of each trail within the IMP-L, IMP-C and NORM (top).

### Selected features

Features selected generally include the ankle, knee, hip, center of mass (CoM) and vertical ground reaction forces (GRF). Only the features selected in DLDJ, SLDJ, DLCMJ, SLCMJ and SLHop are reported, as HuHo, CoDP and CoDU have demonstrated a clear need for a larger number of features to perform well. Throughout this paragraph, the phrase: 10 to 16 %; corresponds to percentage of the movement cycle not to classification accuracy.

Features selected in the DLDJ^3^ can be linked to jump height (selection of CoM vertical velocity prior to take off) and symmetries during the latter part of the first ground contact.

Features selected in the SLDJ^4^ can be linked to jump height (CoM vertical velocity prior to take off), knee and hip kinematics during the middle and latter part of the first ground contact as well as a possible cheating pattern - as evident by the selection of a resultant CoM velocity at impact. This feature should not hold any meaningful information towards classification as every trial was recorded from the same drop height. As such, impact CoM velocity should theoretically be nearly the same across all trails. However, results demonstrate that some information is held in this feature and the ?unusual? error pattern within the SLDJ. It is possible that trials of the class IMP-L might have ?changed? the drop height by adapting a stepping down pattern that has been undiscovered during the biomechanical assessment even with frequent lab quality assessments.

Features selected in the DLCMJ^5^ can be linked to performance (resultant CoM velocity prior to take off) and symmetries in knee rotation and hip angular velocity and the interconnection of segments. While symmetry of knee rotation is a commonly reported feature 35?37 the feature symmetry CoM in anterior hip orientation has not been examined by other studies as it has only been introduced recently38. This measure describes the position of the CoM in a joint position dependent orientation and is influenced by foot positioning, the knee joint orientation and the trunk position. The selection of these CoM in joint features highlights the importance of considering the interconnection of joints and segment during a movement. Further, relevance of the interconnection of joints and segment is also demonstrated by the frequent selection of features that describe the CoM within a joint orientation across the examined exercises.

Features selected in the SLHop^6^ can be linked to the ankle-ground interaction at the impact (ankle flexion velocity and GRZ at phases post impact) and knee flexion at the end of the eccentric phase.

### Practical Implications

Return to play after ACL surgery / prevention of subsequent re-injury are not always guaranteed [44, 45] and this might be in part due to absence of clear criteria identifying if an athlete has returned to pre-injury levels or completed rehabilitation. Current clinical testing batteries often utilize biomechanics to assess a movement quality. However, there is little consensus on the appropriateness of biomechanical analysis or and specific exercise tests and measures when differentiating between two specific groups. This study demonstrates that biomechanical data hold enough information to differentiate between IMP-L, IMP-C and NORM with classification accuracies above 75 %, that a large proportion of individuals included within the control group do not represent a normal movement pattern and that the probability of membership to a class (in this case the NORM class) might allow the generation of a “healthy” healthy score. Such a score can give an objective measure of how close a trial is to a desired class and might present a clear criterion if an athlete has returned to normal or has completed rehabilitation. The “healthy” NORM pattern could be represented by trials those were continuously classified as NORM, and might be considered “low risk” for prospective injury. As the risk of a second ACL injury to the same or contralateral limb is considerably higher than risk of ACL injury in previously un-injured healthy subjects41?44 the here presented framework may be able to even judge risk of injury by using the strength to the NORM group from a trial. However, this assumption cannot and was not tested and need confirmation on an prospective data set containing a similar cohort.

Based on the findings, a biomechanical testing protocol should contain a single and double leg-jumping task (SLCMJ, DLCMJ and DLDJ), as they were able to differentiate trials with high accuracy with only a few features. More complex task, HuHo and CoDP should also be included, for individuals in later stage of rehabilitation, as they challenge the athlete’s ability in more than one plane and were also able to differentiate limbs with high accuracy - with a large number of features. The SLDJ and CoDU demonstrated the lowest ability to differentiate limbs due to their complexity and the range of possible execution strategies. However, all exercises demonstrated that they contain valuable information.

Findings highlight problems with the assumption that the majority of a control group demonstrates a healthy pattern, especially with increasing complexity of a task. With increasing complexity of the exercises the percentage of ?true? members of the NORM class decreased. This could be a reason for conflicting findings in studies. Only the SLCMJ demonstrated a large homogeneity within the control group, while other findings of other exercises suggested that up to 65 % of the limbs within NORM did not present a true NORM pattern and hold some characteristics of IMP classes. The introduced framework can identify true class members through repeated classification. Classification approaches have been applied previously and have demonstrated the ability to enhance the insight into movement data 11,26,32.

### Limitations

Four limitations exist in this study. The first limitation of this study is that the tested machine learning techniques have not been optimized - e.g. the k-nearest neighbor might have performed better using a different k. Accuracies observed are very likely be higher with an ?fine tuned? machine learning technique. The second limitation is the definition of key phases and the simulation to detect key phases. In some cases, a waveform, e.g. angular velocity, can be multimodal (multiple local maxima’s) and extraction the maximal and minimal values as well as their position might have improved classification accuracies, while the detection of key phases through the simulation might also have lowered the classification performance as some important phases might not have meet the subjectively chosen criteria. Thirdly, this study has only used the best trial of an individual and this may have introduced a maximal effort bias. Lastly, a decreased conditioning level within the IMP classes might have improved classification performances.

## Conclusion

This study introduced and tested a framework that combined an automatic feature extraction with machine learning and assessed its ability to differentiate a movements performed by limbs with ACL reconstruction (IMP-L), limbs contralateral from IMP-L (IMP-C) and limbs of a control group (NORM). Findings of this study demonstrate that predictor features extracted from biomechanical data hold valuable information for assessing rehabilitation progress/status, highlighting the potential of movement analysis and machine learning, that a large portion of a control group might not, in identifying injury risk and rehabilitation status. Overall, biomechanical data requires advanced statistics to identify true representations of a group movement pattern, which suggests that probabilities to previously identified patterns may be appropriate to objectively judge injury risk and rehabilitation status.

## Acknowledgments

We are thankful to our colleagues who provided expertise that greatly assisted the data processing or though process of this study.

## Footnotes

1 For the DLCMJ, SLCMJ, DLDJ, SLDJ, HuHo, CoDP and CoDU the end of the eccentric phase was moved to 65, 61,45, 43, 50, 44 and 45 % of the movement cycle, respectively. For the SLHop the peak negative power was moved to 25 % of the movement cycle.

2 Measure included were: GRF (x, y, z), GRF impulse (x, y, z), CoM velocity (xy, xyz, z), CoM Power (x, y, z), CoM in pelvis (x, y, z), CoM in hip (x, y, z), CoM in knee (x, y, z), CoM in ankle (x, y, z), joint angle of ankle, knee, hip, pelvis, thorax and thorax on pelvis in sagittal, frontal and ransversal planes, joint angular velocities of ankle, knee, hip, pelvis, thorax and thorax on pelvis in sagittal, frontal and transversal plane, joint powers, moments, work and impulse of ankle, knee, hip and pelvis in sagittal, frontal and transversal plane, time and foot angle on pelvis.

3 Selected were: symmetry GRF (81 to 86 %), CoM vertical velocity (79 to 83 %), symmetry knee flexion angular velocity (75 to 80 %) and symmetry CoM in anterior ankle orientation (1 to 5 %).

4 Selected were: CoM vertical velocity (94 to 101 %), resultant CoM velocity (1 to 6 %), CoM in anterior knee orientation (57 to 69 %), hip abduction angle (72 to 79 %) and knee abduction angle (95 to 101%).

5 Selected were: symmetry knee rotation angle (29 to 33 %), resultant CoM velocity (87 to 94 %), symmetry CoM in anterior hip orientation (57 to 69 %), hip flexion angular velocity (72 to 79 %).

6 Selected were: Ankle flexion angular velocity (7 to 11 %), knee flexion angular velocity (94 to 101 %), GRF (14 to 18 %) and CoM in anterior knee orientation (33 to 42 %).

